# *Wolbachia* interacts with the microbiome to shape fitness-associated traits during seasonal adaptation in *Drosophila melanogaster*

**DOI:** 10.1101/2022.05.31.494239

**Authors:** Lucas P. Henry, Michael Fernandez, Scott Wolf, Julien F. Ayroles

## Abstract

The microbiome contributes to many different host traits, but its role in host adaptation remains enigmatic. The fitness benefits of the microbiome often depend on ecological conditions, but fluctuations in both the microbiome and environment modulate these fitness benefits. Moreover, vertically transmitted bacteria might constrain the ability of both the microbiome and host to respond to changing environments. *Drosophila melanogaster* provides an excellent system to investigate the evolutionary effects of interactions between the microbiome and the environment. To address this question, we created field mesocosms of *D. melanogaster* undergoing seasonal adaptation with and without the vertically transmitted bacteria, *Wolbachia pipientis.* Sampling temporal patterns in the microbiome revealed that *Wolbachia* constrained microbial diversity. Furthermore, interactions between *Wolbachia* and the microbiome contributed to fitness-associated traits. *Wolbachia* often exerted negative fitness effects on hosts, and the microbiome modulated these effects. Our work supports recent theoretical advances suggesting that hosts in temporally fluctuating environments benefit from flexible microbial associations with low transmission fidelity—specifically when changes in the microbiome can better enable host phenotypes to match environment change. We conclude by exploring the consequences of complex interactions between *Wolbachia* and the microbiome for our understanding of eco-evolutionary processes and the utility of *Wolbachia* in combating vector-borne disease.

## INTRODUCTION

The microbiome shapes many different traits in many different eukaryotic hosts, contributing to behavioral, metabolic, and immunological phenotypes (1–3). While progress has been made in identifying the functional effects of the microbiome in laboratory settings, the eco-evolutionary forces that generated the links between host and microbiome remain poorly understood (3, 4). One key reason is that the phenotypic effects and the potential fitness benefits of the microbiome on their host often depend on the local environment (3, 5); changing environments can shift the relative costs and benefits of host-microbe interactions. Furthermore, the microbiome itself is dynamic. Feedback between the host and the environment can also change the composition and function of the microbiome (6, 7). The dynamic nature of the microbiome may itself be a key feature of host-microbiome interactions, contributing to buffering the effects of environmental stress and potentially conferring key adaptive benefits for the host.

Transmission fidelity can also influence the evolutionary importance of the microbiome (3, 8, 9). Transmission fidelity refers to how faithfully the microbiome is shared across generations, between parents and offspring. Generally, for the microbiome to influence host fitness, microbes benefit their hosts, and hosts faithfully transmit the beneficial microbes to the next generation (3, 8). Hosts tend to evolve strict control of microbial transmission through vertical transmission to maintain these beneficial interactions (9), such as the intricate molecular mechanisms that govern classic symbioses, like the *aphid-Buchnera* association (10). However, strict control can limit the acquisition of other potentially more beneficial microbes, constraining hosts and microbes to the ecological conditions that generated the associations in the first place (3, 11, 12). Recent theoretical advances suggest that for organisms that occupy habitats with variable environments (e.g., seasonality or anthropogenic change), lower transmission fidelity and increased flexibility in the microbiome through environmental acquisition may actually benefit hosts (9). If the microbiome provides functions that benefit hosts, then flexibility in the microbiome may help hosts better match phenotypes to changing environments. Notably, the flexibility in the microbiome may also depend on microbe-microbe interactions. Vertically transmitted microbes are often present at embryogenesis, while other environmentally acquired microbes colonize throughout different points, over development and throughout the lifespan of hosts (13). Priority effects by the vertically transmitted microbes may thus facilitate or impede variation in the environmentally acquired microbiome (14, 15), but the fitness effects of priority effects on hosts remain poorly characterized.

Understanding these fundamental processes may also provide novel solutions to pressing problems in public health (e.g., vector-borne diseases) and agriculture (e.g., pesticide resistance). A particularly notable vertically transmitted bacterium that is currently being developed to suppress vector-borne disease is *Wolbachia. Wolbachia* are intracellular alpha-proteobacteria, vertically transmitted, and found in an estimated 40-60% of all arthropod species (16, 17). *Wolbachia,* classically known for reproductive manipulations (18, 19), also impacts other life history, metabolic, and immunological traits (20, 21). One key trait is pathogen blocking which occurs when *Wolbachia* impedes the establishment of pathogens in the host, including pathogens vectored by mosquitoes (22, 23). Indeed, *Wolbachia* infected mosquitoes can reduce the incidence of dengue by ~70% in some locations (24). Yet, introducing *Wolbachia-infected* mosquitoes is labor intensive as *Wolbachia* does not naturally infect most of the mosquitoes that transmit diseases, requiring the rearing of millions of mosquitoes to spread *Wolbachia* effectively enough to provide protection against vector-borne diseases (25).

To increase the efficiency of *Wolbachia* introductions, a better understanding of how *Wolbachia* shapes host fitness is needed. The fitness costs and benefits of *Wolbachia* often depend on ecological conditions (21, 26, 27). With climate change potentially reshaping the distribution of vector-borne pathogens (28), identifying new mechanisms that buffer environmental stressors will be needed. The adaptive potential of the microbiome provides one path forward, as rapid evolution in the microbiome may facilitate local adaptation (3). However, much about the evolutionary impacts of the interactions between hosts, *Wolbachia,* and the microbiome remain cryptic (29–31).

Elucidating the eco-evolutionary processes that shape the fitness effects of *Wolbachia* has long been facilitated by research in *D. melanogaster. Wolbachia* is prevalent in *D. melanogaster,* infecting ~30% of the *Drosophila* Stock Center (32) as well as many natural populations (33, 34). Pathogen blocking by *Wolbachia* was first discovered in *D. melanogaster* (35, 36), and mechanisms involved in pathogen blocking discovered in *Drosophila* often apply to mosquitoes (22, 37, 38). Additionally, the *Drosophila* microbiome is relatively simple (<20 species), environmentally acquired, and shapes many different traits (39). Previous research suggests that *Wolbachia* has conflicting effects on the microbiome, with both antagonistic (40) and beneficial (41) effects on *Acetobacter* and *Lactobacillus,* the dominant bacteria in the fly microbiome. These bacteria are also implicated in shaping adaptation in *Drosophila.* In a field mesocosm experiment, inoculation with *Acetobacter* and *Lactobacillus* rapidly generated genomic divergence within five generations during fly adaptation to a seasonally changing environment (42); however, all flies were infected with *Wolbachia.* In the laboratory, a metaanalysis of experimental evolution in *Drosophila* found that *Wolbachia* and microbial diversity frequently responded to artificial selection (43), suggesting that the interactions between *Wolbachia* and the microbiome may contribute to host evolution. Thus, *D. melanogaster* is an excellent model to study the evolutionary interplay between host, microbiome, and the environment.

Here, using field mesocosms, we performed longitudinal sampling to study the microbiome dynamics in *D. melanogaster* with and without *Wolbachia* during adaptation to a seasonally changing environment (Fig. 1). If bacteria with high transmission fidelity (i.e., *Wolbachia)* shape the microbiome and fitness effects on the host, then *Wolbachia* infected (W+) flies may differ in seasonal adaptation compared to *Wolbachia-free* (W-) flies. We combined our longitudinal microbiome dynamics with phenotyping for fitness-associated traits to understand how *Wolbachia* and microbiome interactions shape host adaptation.

**Figure 1:**
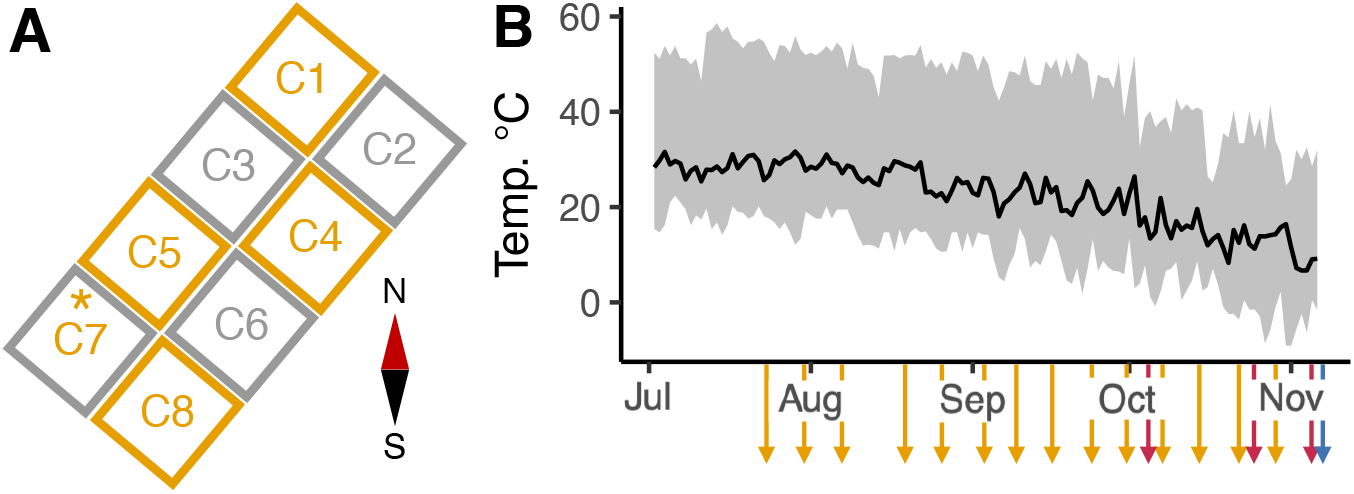
Experimental design. A) Layout of the eight cages, with compass showing North-South orientation. Cages are colored by *Wolbachia* status, with gray representing *Wolbachia*-free (W-) and orange representing *Wolbachia* (W+) flies. All cages maintained the initial *Wolbachia* status, except for C7, which converted to *Wolbachia* infected in mid-August (~Day 57). B) Temperature and sampling regime over the season. Cages were seeded with flies on July 2, 2019 and ended November 6, 2019. Daily mean temperature is shown in black line and range shown in gray. The colored arrows represent sampling points. Microbiome was sampled weekly (orange, N=14 timepoints) beginning Day 24 (third week of July) until Day 120. Flies were periodically phenotyped for starvation resistance as a proxy for fitness (red, N=3 timepoints). Finally, at the end of the experiment (blue), longevity and fecundity were measured in females to test if *Wolbachia* status and microbiome variation influenced seasonal evolution.

## RESULTS

### Season shapes the composition of the microbiome

Flies were sampled weekly over the season beginning three weeks after the experiment started in July 2019. Fly populations maintained their *Wolbachia* status throughout the experiment, except for Cage 7. Cage 7 was initially *Wolbachia*-free, but we detected *Wolbachia* on the Day 57 sampling and subsequently at the rest of the sampling points. We note that while *Wolbachia* is part of the microbiome, for simplicity, we will refer to the microbiome as the bacterial community that primarily infects the gut but can also survive outside of the host (39).

#### The microbiome was predominantly composed of four bacteria

*Acetobacter, Commensalibacter, Providencia*, and *Wautersiella* (Fig. 2). At the start of the season, *Acetobacter* and *Providencia* were the dominant bacteria. *Acetobacter* peaked early in the season (Day 57), and then was replaced by another bacteria in the Acetobacteraceae family, *Commensalibacter,* which dominated for the rest of the season. *Providencia* fluctuated, with peaks at the beginning and end. *Wautersiella* peaked in the middle of the season (Day 85), but was largely absent at the beginning and end. W- and W+ flies generally harbored the same bacteria, but the relative abundance differed over the season in complex ways; differences between *Wolbachia* status varied from one time point to the next.

**Figure 2:**
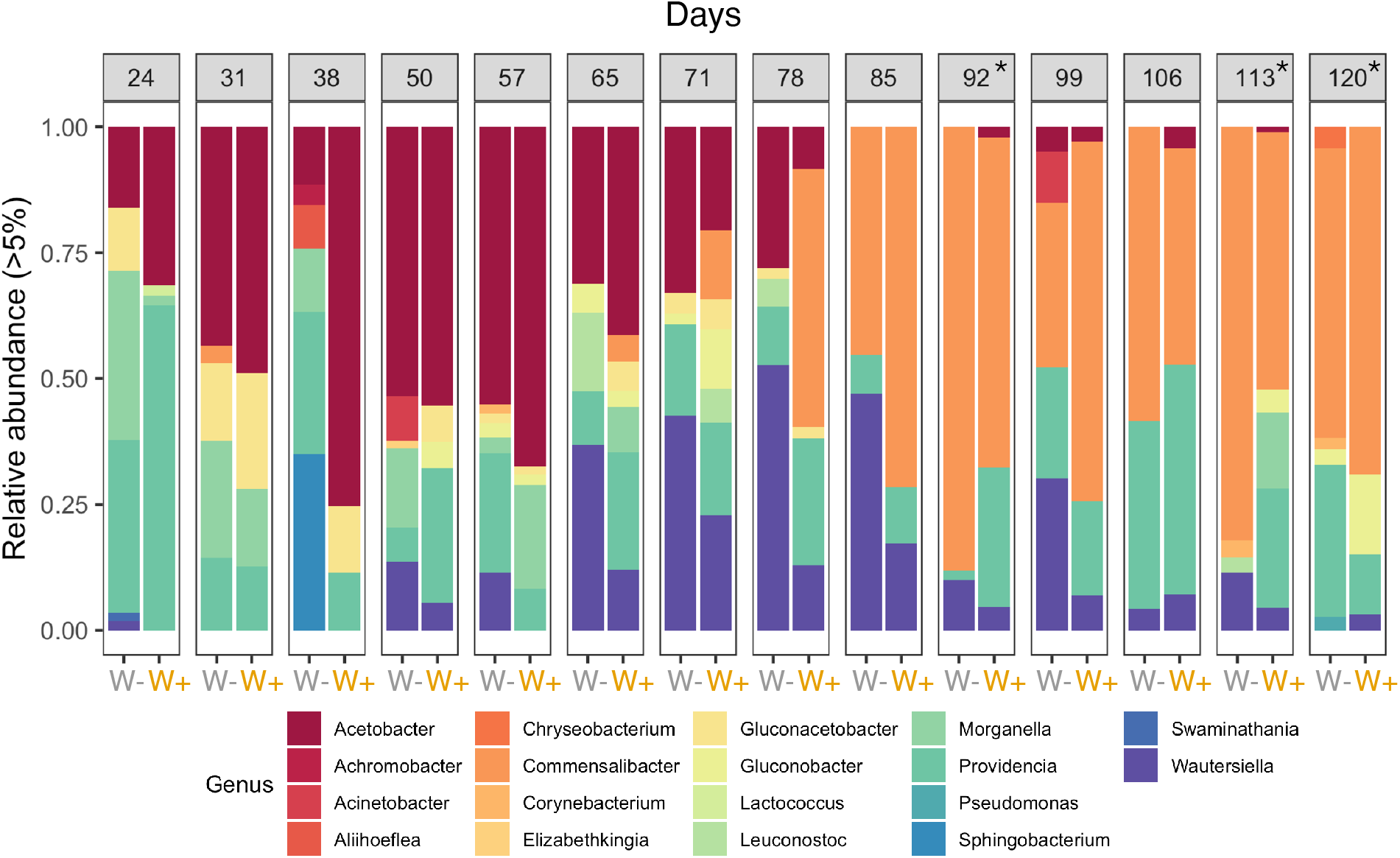
Microbiome composition over growing season. Groups are faceted by sampling date with averaged *Wolbachia-free* (W-) and *Wolbachia* (W+) populations. Asterisks denote the dates that were paired with the fitness-associated phenotyping later in the season. Colors represent the different genera. *Acetobacter* was replaced by *Commensalibacter* by the end of the season, while *Wautersiella* peaks in the middle. *Providencia* was intermittently present throughout the growing season, though most abundant at the beginning and end of the season.

### *Wolbachia* constrains microbiome diversity in seasonally changing environment

If vertically transmitted microbes constrain the ability of the microbiome to respond to environmental fluctuations, then *Wolbachia* infection could reduce microbial diversity in two ways. First, *Wolbachia* infection could reduce the complexity of the community within a population (i.e., alpha-diversity). Second, *Wolbachia* infection may change how community turnover proceeds over the season (i.e., beta-diversity). Through longitudinal sampling across replicated W+/W− populations, we assessed how *Wolbachia* infection shaped microbiome dynamics.

Indeed, W+ flies exhibited reduced fluctuations in alpha-diversity (Fig. 3). Shannon diversity fluctuated throughout the season, while phylogenetic diversity stabilized towards the end of the growing season (Fig. 3A). W- flies accumulated significantly more changes to Shannon diversity than W+ flies (Fig. 3B, Kruskal-Wallis X^2^ = 5.00, df = 1, p = 0.03). However, while a similar trend was observed for phylogenetic diversity, it was not statistically significant (Kruskal-Wallis X^2^ = 2.69, df = 1, p = 0.10).

**Figure 3:**
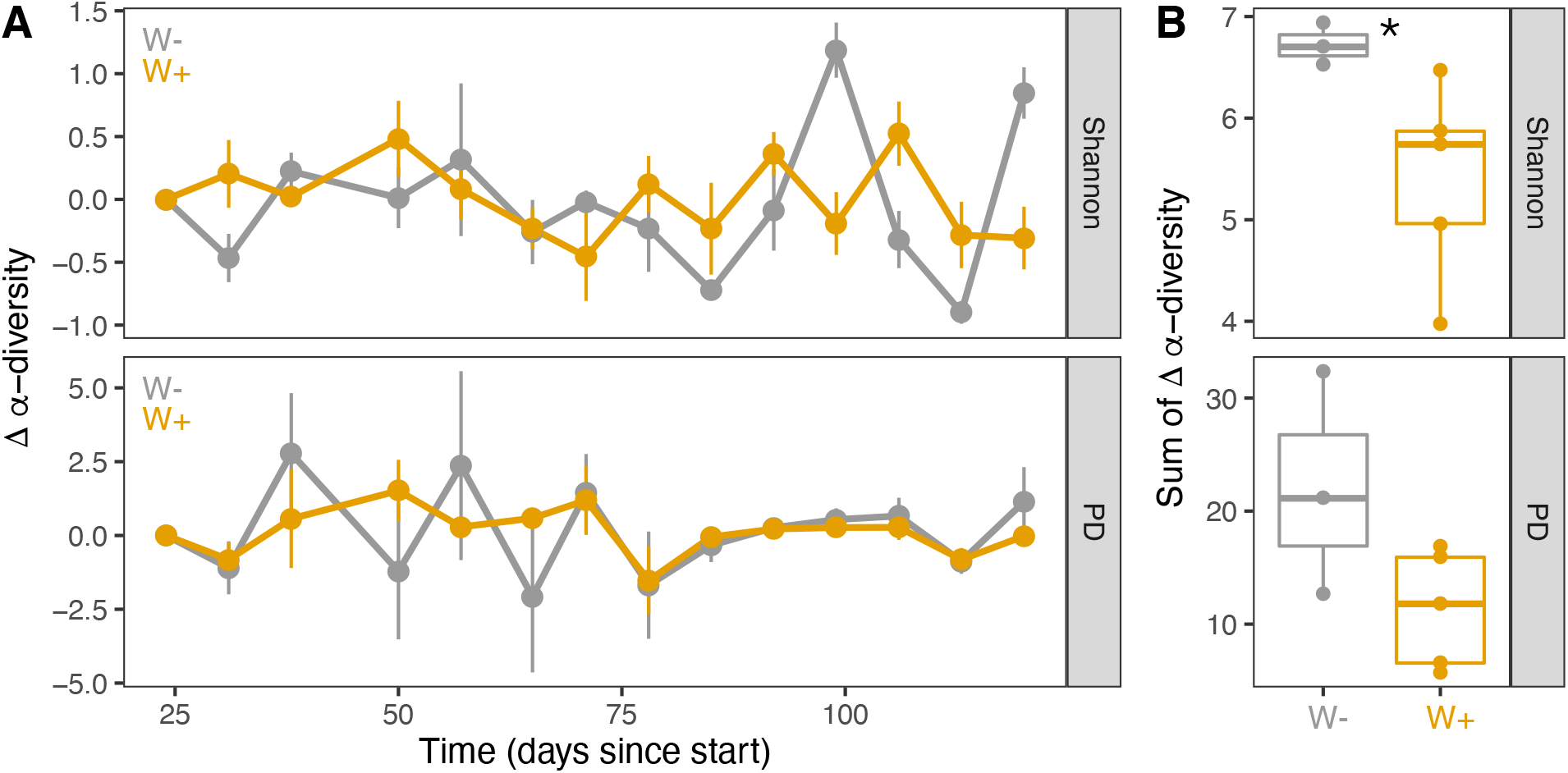
Alpha diversity fluctuates more in *Wolbachia-free* populations over the season. A) Change in two alpha-diversity measures (Shannon diversity; PD = phylogenetic diversity) colored by *Wolbachia* status. Lines represent the mean change in alpha diversity and error bars are standard error. For Shannon diversity, fluctuations increased at the end of the season, particularly for W- flies. For PD, both populations fluctuate more at the beginning of the growing season. B) Summation of the changes in alpha diversity over the season by each cage. While PD tended to be greater for W-, only the change in Shannon diversity for W-populations was significantly higher than W+ flies (Kruskal-Wallis X^2^ = 5.00, df=1, p=0.03).

Community turnover was also shaped by *Wolbachia* infection status (Fig. 4). Principal coordinate analysis using Bray-Curtis dissimilarity (BC) showed that time (i.e., days since the start of experiment) significantly shaped differences between microbiomes (Fig. 4A, PERMANOVA: R^2^ = 0.20, p = 0.001, Supp. Table R1). *Wolbachia* exerted significant, but marginal effects on the microbiome (PERMANOVA: R^2^ = 0.02, p = 0.001, Supp. Table R1). A similar trend was observed using Unifrac distance (Supp. Fig. R1, Supp. Table R2). We illustrated community turnover by showing the temporal trends in the top four abundant bacteria (Fig. 4B). *Acetobacter* and *Providencia* declined first, followed by the peak for *Wautersiella,* then *Comensalibacter* and *Providencia* increased at the end of the season.

**Figure 4:**
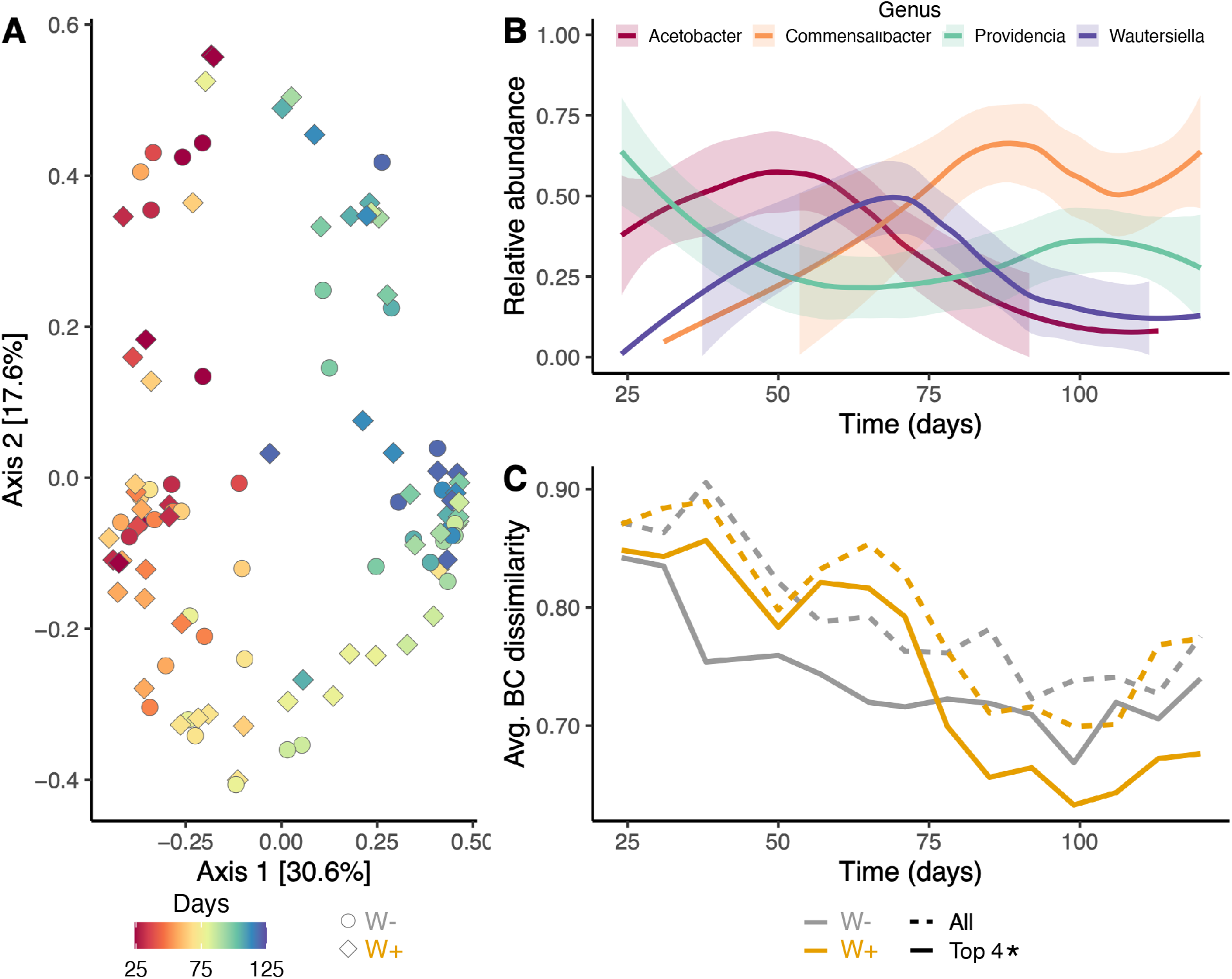
Community composition turnover changes over growing season and interacts with *Wolbachia* status. A)PCoA plot using Bray-Curtis dissimilarity. Color represents time (days since the start), where warm colors represent the summer beginning and cool represent the fall ending. Stape shows *Wolbachia* status. Seasonality significantly affected differences setwee ⋂ all individuals (Supp. Table R1). *Wolbachia* infection exerted significant, but marginal effects. There was no significant! nteraction between *Wolbachia* and seasonality. B) Temporal dynamics in the four most abundant genera illustrate community turnover. Lines represent the average across all cages for each genus with loess smoothing, and 95% confidence intervals are shaded. C) Community turnover over the growing season. Lines represent the Bray-Curtis (BC) dissimilarity within each cage, averaged by *Wolbachia* status. Dashed lines show BC for the complete microbiome, while the solid lines show for the top four most abundant genera visualized in B. BC dissimilarity decreases over the growing season. For the top four abundant genera, *Wolbachia* interacted significantly with community turnover, where *Wolbachia* populations were initially more dissimilar, but by the end of the growing season, became more similar than *Wolbachia-free* populations (Supp. Table R3).

We next assessed the mean BC for each population cage over the growing period. High BC is associated with more community turnover, while lower BC values correspond to greater similarity. In general, BC decreased over the season (Fig. 4C). Notably, *Wolbachia* interacted with seasonality to shape community turnover for the four dominant bacteria *(Wolbachia* x time interaction: Wald X^2^ = 12.91, df = 1, p = 0.0003, Supp. Table R3). W+ populations were initially more dissimilar than W-, but became more similar by the end of the season. However, the *Wolbachia* x time interaction was not detected for the complete community *(Wolbachia* x time interaction: Wald X^2^ = 1.01, df = 1, p=0.31, Supp. Table R4).

Through reducing microbiome diversity and turnover, *Wolbachia* infection likely constrained the ability of the microbiome to respond to the seasonally changing environment. So far, we have considered only microbe-microbe interactions. However, these microbial communities are also changing in the context of the host response to the changing environment.

### Interactions between *Wolbachia* and microbiome shape fitness-associated host traits

If interactions between *Wolbachia* and the microbiome influence how the host responds to changing environments, then we would expect to see differences in fitness-associated phenotypes emerge over the course of the experiment. To test this, we performed periodic phenotyping towards the end of the season for starvation resistance. Starvation resistance reflects the nutritional reserves used in both reproduction and survival in challenging environments. We phenotyped flies at three time points paired with longitudinal microbiome profiling to assess how interactions between *Wolbachia,* microbiome, and the changing environments shape the response in the fly populations.

Starvation resistance varied over the season (Fig. 5A). For the microbiome, we focused on the effects of the most frequent bacterium found at the three time points, *Commensalibacter.* For the first time point (Day 96), only sex affected starvation resistance; males starved twice as fast as females (β = 2.08 ± 0.20 SE, p < 0.0001, Supp. Table R5). At the next time point (Day 116), increased *Commensalibacter* relative abundance was associated with decreased starvation resistance ( β = 1.21 ± 0.38 SE, p = 0.001, Supp. Table R6). W+ flies starved approximately twice as fast as W- flies ( β = 0.52 ± 0.18 SE, p = 0.003). There were moderate *Wolbachia* x sex interactions, but this effect was not statistically significant and was removed from the model. However, at the final time point, *Commensalibacter* and *Wolbachia* x sex interactions significantly influenced starvation resistance (Supp. Table R7). Again, *Commensalibacter* relative abundance negatively affected starvation resistance ( β = 1.33 ± 0.48 SE, p = 0.006). Sex-by-*Wolbachia* interactions also emerged ( β = −0.71 ± 0.30 SE, p = 0.017). While W-females had greater starvation resistance than males, there was no difference for W+ flies, resulting in a substantial reduction in fitness for females especially. Overall, *Wolbachia* and the microbiome shaped starvation resistance, particularly at the end of the season (Fig. 5B).

**Figure 5:**
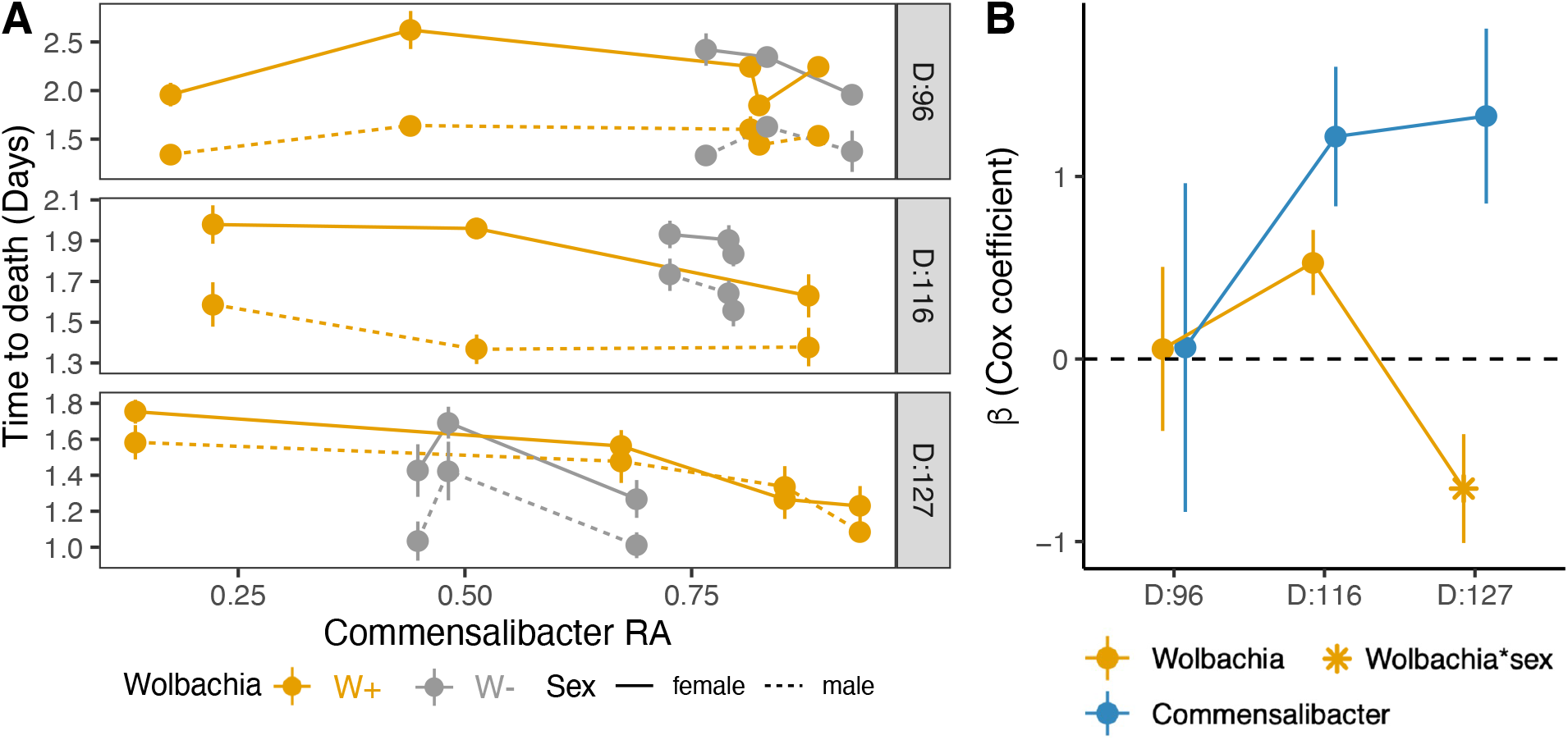
*Wolbachia* interacts with *Commensalibacter* to shape starvation resistance. A) Points represent the mean time to death (days) for each cage with standard error. X-axis shows the relative abundance of *Commensalibacter.* Color represents *Wolbachia,* with solid lines for females and dotted lines for males. Facets show the three time points collected over the later end of the growing season, labeled by the date of collection. For all time points, sex affected starvation resistance, with males starving more quickly than females. Earlier in the growing season at Day 96 (N: W+ = 107, W- = 77), neither *Wolbachia* nor *Commensalibacter* affected starvation time. At Day 116 (N: W+ = 93, W- = 92), *Commensalibacter* was negatively associated with starvation resistance (Supp. Table R6). Furthermore, *Wolbachia* flies were less starvation resistant than *Wolbachia-free* flies (Supp. Table R6). In the final sampling at Day 127 (N: W+ = 110, W- = 86), increased *Commensalibacter* significantly reduced starvation resistance (Supp. Table R7). *Wolbachia* and sex interacted to shape starvation, where *Wolbachia-free* females had higher resistance than males, but there was no difference between sexes for *Wolbachia* flies (Supp. Table R7). B) Model estimates (β) ± standard error from Cox hazard models over the three sampling points that summarizes data shown in Fig. 5A. *Wolbachia* effects are displayed in orange and blue shows the effects of *Commensalibacter.* For D:127, shown are only the model estimates for statistically significant *Wolbachia* x sex interaction. Overall, the effects of *Wolbachia* and *Commensalibacter* increased at the end of the growing season.

Finally, at the end of the season (Day 127), we measured fecundity and lifespan in individual females from each population. Fecundity was measured as the number of pupae that emerged from individual females. Fecundity was zero-skewed, with 50% W+ and 35% W- females producing no pupae (Fig 6A). For the females with non-zero fecundity, pupae produced ranged from a single pupa to 118 pupae. Lifespan also varied (Fig. 6B). While 30% of W+ and 24% of W- females died within the first sampling point (11 days after the egg lay; after Day 11, checked every 4-5 days until all died), lifespan ranged from 12 to 82 days. For the microbiome, we examined the ratio of the two most frequent bacteria observed at the end of the season, *Providencia* (6 populations) and *Commensalibacter* (all 8 populations). There was no significant effect on *Wolbachia* on the *Providencia:Commensalibacter* (Prov:Comm) ratio (Kruskal-Wallis X^2^ = 0.81, df = 1, p = 0.37); though we note the range in W- populations spanned from 0 to 1.2, while W+ populations only from 0 to 0.5 (Fig. 6C).

**Figure 6:**
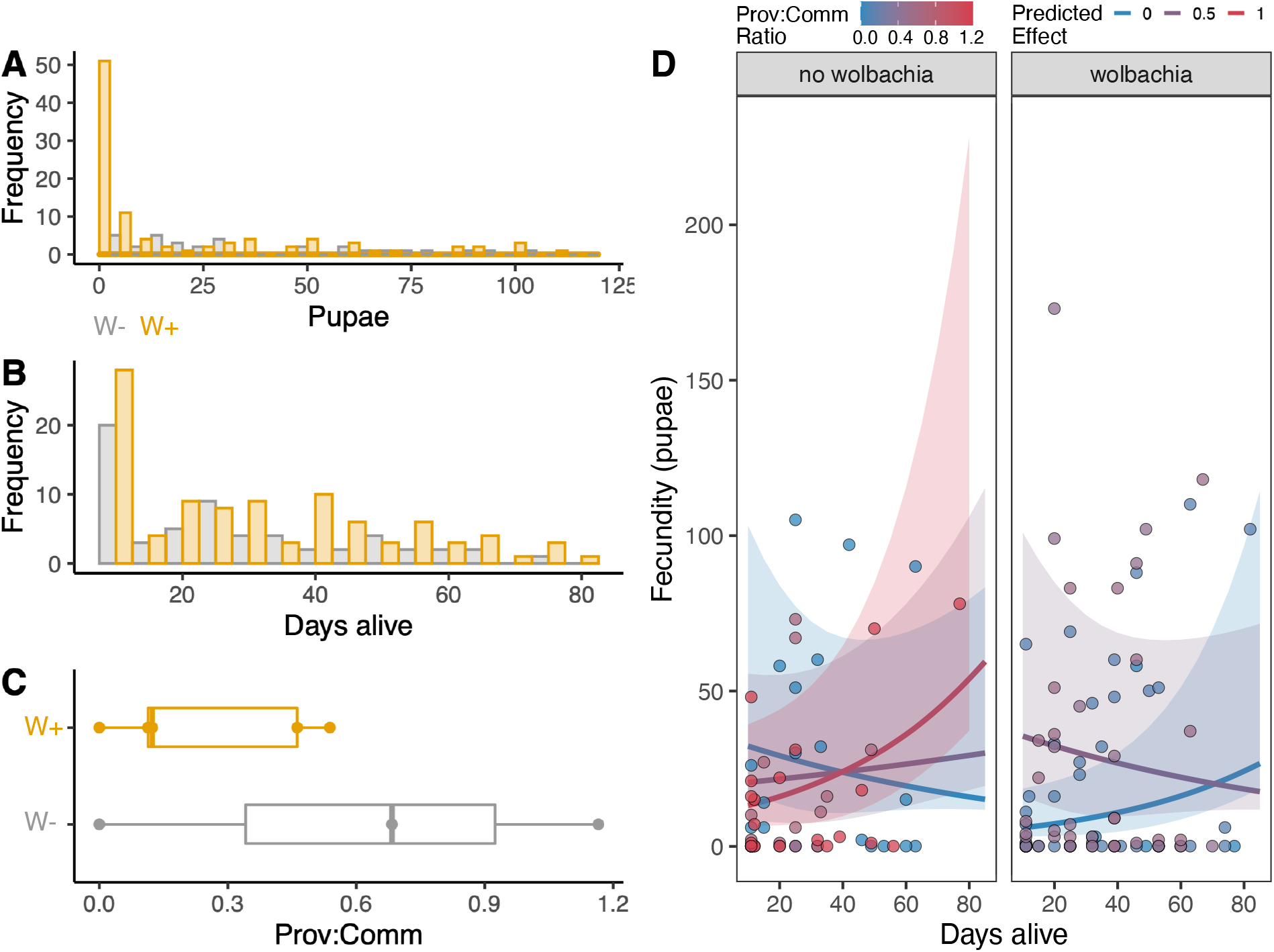
Interactions between *Wolbachia* and the microbiome shape fitness-associated traits following seasonal evolution. Female flies (N=~20/cage, total W+ = 99, W- = 60) were collected at the end of the growing season and individually phenotyped for fecundity (pupae produced) and lifespan (days alive in the lab). A) Distribution of pupae produced from single females, colored by *Wolbachia* status. B) Distribution of lifespan in single females, colored by *Wolbachia* status. C) Box plots showing differences in the ratio of *Providencia* to *Commensalibacter* (Prov:Comm) between *Wolbachia* status. D) Fecundity from the interaction model between lifespan, *Wolbachia,* and microbiome. Prov:Comm ratios are from the final microbiome sampling point, represent averages per each cage, and were modeled at the three Prov:comm levels that captured most of the variation across cages. Lines show the predicted interactions between lifespan, fecundity, and Prov:Comm ratios, faceted by *Wolbachia* status. Points show the measured values and are shaded by the Prov:Comm ratio. The three way interaction significantly predicted fecundity through interactions between lifespan, *Wolbachia,* and microbiome (Supp. Table R8). *Wolbachia-free* flies with high Prov:Comm ratio were less fecund and had shorter lifespans, but *Wolbachia* flies were more fecund with shorter lifespans at low Prov:Comm ratios.

To investigate the effect of the microbiome on the relationship between fecundity and lifespan, we modeled how interactions between *Wolbachia* and microbiome affected the relationship between fecundity and lifespan (Fig. 6D). Broadly, the flies that lived longer were also more fecund. However, the interaction between *Wolbachia* and Prov:Comm ratio shaped this pattern ( β = 0.044 ± 0.02 SE, p=0.01, Supp. Table R8). Primarily, high Prov:Comm ratios were absent in W+ flies, but tended to be associated with higher fecundity in W- flies. More so, W+ flies with low *Providencia* (i.e., Prov:Comm ratio ~ 0) experienced lower fecundity but longer lifespans, while higher Prov:Comm ratio flies experienced higher fecundity but shorter lifespans. However, the effects of the Prov:Comm ratio were minimal on W- flies, though trended towards longer lifespans and higher fecundity with high Prov: Comm ratios. Overall, the interaction between *Wolbachia* and microbiome shaped the fitness of flies following seasonal evolution.

## DISCUSSION

Here, we examined how interactions between vertically transmitted and environmentally acquired bacteria shape adaptation to seasonally changing environments in *Drosophila melanogaster.* The analysis of temporal patterns in the fly microbiome (Fig. 2) suggest that *Wolbachia* constrained microbial diversity (Figs. 3, 4). Furthermore, interactions between *Wolbachia* and the microbiome also contributed to changes in fitness-associated traits. *Wolbachia* often reduced starvation resistance, fecundity and lifespan, but this was mediated by an interaction with two dominant bacteria, *Commensalibacter* and *Providencia* (Figs. 5, 6). Next, we discuss how our results provide insights into the influence of complex interactions between *Wolbachia* and the microbiome on host seasonal evolution.

### *Wolbachia* constraint on flexibility in the environmentally acquired microbiome

W+ flies experienced reduced fluctuations for Shannon diversity compared to W- flies (Fig. 3). We detected a similar trend for phylogenetic diversity, but because the communities are composed of similar taxa (primarily Acetobacteraceae family), the effects of *Wolbachia* on phylogenetic diversity were not statistically significant. Nevertheless, the reduction in Shannon diversity suggests that *Wolbachia* is limiting community complexity. Additionally, *Wolbachia* also reduced community turnover as measured by decreasing Bray-Curtis dissimilarity (Fig. 4C). The four most abundant bacteria responded strongly to *Wolbachia* compared to the total community, suggesting that dominant bacteria potentially regulate the interaction between *Wolbachia* and the microbiome. Together, this suggests the presence of *Wolbachia* changes the capacity for microbial change in response to the seasonally changing environment.

The mechanisms underlying *Wolbachia* interactions with other bacteria are poorly understood. *Wolbachia* has been shown to have conflicting effects on Acetobacteraceae, either suppressing (40) or increasing (41) its abundance. The interaction is likely regulated through indirect mechanisms as *Wolbachia* do not infect the lumen cells where most other bacteria reside (40). *Wolbachia* often interacts with the immune system when protecting against viral pathogens (36, 44), but has limited effects on protection against bacterial pathogens (40, 45, 46). This suggests that the immune system is not directly involved in regulating the interaction between *Wolbachia* and the environmentally acquired microbiome. Temperature may however contribute to mediating the temporal dynamics of different bacteria. *Wolbachia,* like other intracellular bacteria, are thermally sensitive, with extreme temperatures exerting negative effects on intracellular bacteria and host fitness (47, 48). In *Drosophila,* both high (>28°C) and low (<20°C) temperatures decrease *Wolbachia* abundance and phenotypic effects (e.g., pathogen blocking and reproductive manipulations) in laboratory settings (34, 49, 50). However, bacteria from the fly microbiome are commonly cultured at 30°C (51), with *Providencia* as high as 37°C (46), suggesting these bacteria are more tolerant of temperature variation. During our experiment, mean daily temperature ranged from 26.6°C to 6.6°C, with temperatures as extreme as 55.9°C in full sun and −3.6°C observed (Fig. 1B). Differences in microbiome diversity may result from differential growth across bacterial species and the consequences of *Wolbachia* sensitivity to extreme temperatures. Taken together, our results suggest that environmental variation can alter host-microbe interactions through complex responses to abiotic (e.g., temperature) and biotic factors (e.g., microbe-microbe interactions).

While more work is necessary to identify the specific mechanisms underlying *Wolbachia-* microbiome interactions, our results highlight how *Wolbachia* infection constrains microbial diversity. The constraint on microbial diversity may benefit hosts by linking beneficial microbes to phenotypes that buffer specific environmental stressors and limit potentially negative interactions with deleterious microbes (3). However, too much constraint on the microbiome may cost host fitness if rapid microbial change can provide novel solutions to new selective pressures experienced in fluctuating environments ( 3, 9). Both of these predictions require a better understanding of how interactions between *Wolbachia* and the environmentally acquired microbiome shape host phenotypes.

### *Wolbachia* and microbiome interact to shape fitness-associated traits in hosts

For the microbiome to influence host evolution, host phenotypes should change in response to microbial variation (3). In *Drosophila,* both *Wolbachia* and the environmentally acquired microbiome often shape variation for a wide range of phenotypes (39, 52–55). By examining changes in fitness-associated traits over the course of the season, we identified shifts in the relative importance of interactions between *Wolbachia* and the microbiome for host adaptation.

Both *Commensalibacter* and *Wolbachia* impacted starvation resistance, but the effects depended on the sampling time point (Fig. 5). At the first time point (Day 96), neither had an effect. *Commensalibacter* had direct effects on starvation resistance on both subsequent time points; increased *Commensalibacter* abundance was associated with decreased starvation resistance. Starvation resistance in flies is predominantly determined by the amount of lipids stored (56, 57), and while the effects of *Commensalibacter* on lipid stores are unknown, many of other bacteria in the Acetobacteraceae family reduce lipid storage (58, 59). Other bacteria, like *Lactobacillus,* are typically associated with increased lipid storage and found in fly populations from colder climates (60); however, we did not detect abundant *Lactobacillus* in our study. *Wolbachia* also affected starvation resistance. W+ flies experienced decreased starvation resistance compared to W- flies. Other studies have not found an effect of *Wolbachia* on starvation resistance in laboratory populations of *Drosophila* (26, 61), but *Wolbachia* effects are often context dependent on both host genotype and ecological conditions (26, 62). The effects of *Wolbachia* varied across the three sampling points, with both *Wolbachia* and *Commensalibacter* having larger effects on starvation at the end of the season (Fig. 5B). Intriguingly, *Wolbachia-by-sex* interactions significantly decreased starvation resistance in the final time point (Day 127), as there was no difference in starvation time between males and females only for W+ flies. Reproductive manipulations by *Wolbachia* can target both males and females, biasing transmission of Wolbachia at the expense of host fitness (20, 63, 64). Our results suggest that W+ females exhibited the greatest reduction in starvation resistance at the end of the season, potentially because the complex interaction between *Wolbachia* and the microbiome reduced the ability of females to store lipids in the seasonally changing environment.

The other fitness-associated traits, fecundity and lifespan, were also shaped by interactions between *Wolbachia* and the microbiome at the end of the season (Fig. 6). Notably, for W+ flies, increased *Providencia* shifted life-history trade-offs, with higher fecundity, but short lifespans. However, when *Providencia* was low or absent, W+ flies lived longer and produced more offspring. Only marginal effects were observed for W- flies. The three-way interaction, combined with the starvation resistance results, points to one potential mediator–the insulin/insulin-like growth factor signaling (IIS) pathway. The IIS pathway helps maintain metabolic homeostasis by shaping the balance between carbohydrate availability and lipid storage, and consequently, life-history tradeoffs in *Drosophila* and many animals (65). The microbiome also modulates expression of several genes within the IIS pathway, including insulin receptors (66). However, not all bacteria contribute in the same way; different *Acetobacter* species modulate the activity of key components of the IIS pathway in different ways (67). *Wolbachia* also increases insulin signaling in *Drosophila* (54). Furthermore, many genes within the IIS pathway are highly pleiotropic, and polymorphisms in alleles within the IIS pathway also contribute to variation in life history traits associated with adaptation to ecological differentiation along a latitudinal cline in *Drosophila* (68). Taken together, complex interactions between *Wolbachia* and the microbiome may shape how hosts allocate nutrition and shift life-history strategies to buffer environmental change.

The complex interactions for starvation resistance, fecundity, and lifespan suggest that *Wolbachia* exerted fitness costs on *Drosophila.* We note that we only assessed fecundity early in life and did not account for how fecundity changed over the lifespan. Nevertheless, our work here contributes to the evidence for complex, context dependence of *Wolbachia* effects on host fitness (26, 27, 53).

### Implications for microbiome interactions in seasonally evolving populations

Here, *Wolbachia* shaped the seasonal changes in the environmentally acquired microbiome, and together, both affected fitness associated traits in the flies. These changes overall highlight the potential for variation in microbe-microbe interactions to shape seasonal evolution in hosts–however, the missing link is whether the microbiome changed the host response to selection (3). To understand if the microbiome buffered or changed the host evolutionary trajectory, genomic sequencing is necessary. Comprehensive genomic analyses of fly, *Wolbachia,* and the microbiome will provide deep insights into evolutionary processes shaping seasonal evolution.

Previous work in *Drosophila* has shown how other microbiome manipulations shaped seasonal evolution. In a study where flies were inoculated with either *Acetobacter* or *Lactobacillus* at the start of the season, the different bacteria drove genomic divergence between fly populations in only five generations (42). *Acetobacter* enriched fly genomes for alleles associated with southern populations, where *Acetobacter* is also more common (60). Similarly, *Lactobacillus* enriched for alleles associated with northern fly populations. The fly populations in these experiments were all infected with *Wolbachia,* but our flies lacked *Lactobacillus,* so applying these findings to our results is only speculative. Nonetheless, taken together, different microbial communities may lead to different evolutionary trajectories. In a sense, the host genome may be tracking the changes in the microbiome. As *Wolbachia* decreases the capacity for change in the microbiome, evolution in the host genome also is likely to change, much like adaptive tracking (9, 69). More work is necessary to understand the linkages between host and microbiome evolution, but adaptive tracking may depend on hostmicrobe interactions (9). Adaptive tracking in *Drosophila* can occur during seasonal evolution (70), and potentially in the many organisms that live in temporally fluctuating environments–if and how the microbiome contributes to adaptive tracking remains an open question.

Incorporating microbiome interactions with *Wolbachia* adds additional complexity to an already complex system. However, our results provide insights into how the microbiome may modulate the fitness effects of *Wolbachia* on their host. Mismatches between microbes and the environment may be exacerbated by *Wolbachia,* such as the negative association between *Commensalibacter* and starvation resistance (Fig. 5). Interventions to supplement the microbiome with better matched microbes may help *Wolbachia-infected* hosts buffer challenging environments, as we observed in the *Drosophila* microbiome (42, 60). As millions of mosquitoes are needed for these *Wolbachia*-mediated controlled efforts (25, 71), even moderate improvements to survival by the microbiome may help substantially increase the efficacy of *Wolbachia* in reducing vector-borne disease.

In conclusion, when the microbiome changes in seasonally changing environments, *Wolbachia* may modulate effects on fitness-associated traits in the host. While many questions remain, this study contributes to a growing body of literature utilizing the rewilding of laboratory model systems to uncover how eco-evolutionary processes in host-microbe interactions (72– 76). Future work that links host, microbiome, and their interactions will provide fundamental insights into host-microbe evolution as well as novel solutions for applied challenges in public health.

## METHODS

### Fly populations

Flies in this experiment were derived from a round-robin crossing design of the Global Diversity lines (77) and maintained at large population size (>10,000) flies for >100 generations before the field experiment. This base population was naturally infected with *Wolbachia* (W+). To generate the *Wolbachia-free* (W-) population, flies were treated with 0.25 mg/ml tetracycline in the diet for two generations. W- flies were maintained for ~10 generations before the beginning of this experiment. During this pre-experimental maintenance phase, all flies were maintained at 25°C with 12 hour light:dark cycles. All flies, in the lab and field, were reared on a diet composed of 10% glucose, 10% yeast, 1.2% agar with 0.04% phosphoric acid and 0.4% propionic acid as preservatives.

To confirm *Wolbachia* status, we amplified two genes: cytochrome oxidase I (COI) in D. melanogaster (78) and 16S rRNA gene from *Wolbachia* (79). The COI gene served as a positive control as all fly samples should always generate an amplicon. Primer sequences can be found in Supp. Table M1.

### Field design and experimental sampling

Full details can be found in the Supplementary Methods.

The field site was located at Princeton University, NJ (40.34°N, 74.64°W). Eight total cages were constructed at the field site (Fig. 1). Cages were 1.2 m x 0.6 m x 0.6m (height x width x depth) and constructed from polyethylene monofilament fabric with 150×150 micron mesh (Greenhouse Megastore IS-NT-99). Each cage held a temperature/humidity data logger (Elitech USA Temlog 20H) placed on shelving units within the cage that held fly food. Depending on the position in the field, some temperature loggers may have been in direct sunlight during the daytime, but *Wolbachia* treatment was alternated to deal with variation in sun exposure (Fig. 1).

Approximately 2500 flies (equal sex ratio) were placed into each cage at the start of the experiment on July 2-3, 2019. We introduced 1000 flies on July 2 and an additional 1500 on July 3. We maintained populations by providing ~300 ml fly food once per week, allowing for flies to feed, lay eggs, and providing a substrate for larvae to develop. As the population grew, we provided food twice a week, with only one loaf pan/week kept with the developing flies to maintain population size. Generations were overlapping, but we estimate that ~10 fly generations occurred during the experiment from July 2 until November 6, 2019.

We allowed fly populations to stabilize for the first three weeks (~1-2 fly generations). Following this, we sampled flies weekly to check *Wolbachia* status and characterize change in the fly microbiome. 10 flies were collected from each cage and PCR confirmed for *Wolbachia* status as previously described and then saved for 16S rRNA amplicon sequencing. This resulted in 14 timepoints over the season.

To understand how *Wolbachia* and microbiome change affected fly fitness, we collected age-matched flies for starvation resistance on three dates: Day 96, Day 116, and Day 127. To age-match flies, we performed a separate egg lay in a cage within the cage. Flies were age-matched to 5-7 days old. We note Day 127 flies were not specifically age matched, but collected from within the larger cage, which reflects the standing variation in traits at the end of the season.

To measure starvation resistance, we recorded flies in 15×6.25 mm (diameter x height) acrylic arenas over 4-5 days until death. Arenas contained 1% agar to provide humidity, but no nutritional value. Cages were randomized across observation plates. Full details of recording parameters are in the supplement. We determined the time of death in 2.5 hour intervals to quantify starvation resistance.

For the Day 127 flies, we also measured fecundity and lifespan from individual females to identify whether these fitness-associated traits varied between *Wolbachia* status. Individual females were placed into fly vials with 6 ml fly food and allowed to lay eggs for ~24 hrs. Fecundity from females was measured as the number of pupae that emerged from the 24 hour egg lay. We then flipped the flies into new vials after 10 days. Then, after 10 days, we flipped every 3-4 days until flies died to determine the lifespan of each individual. Fecundity is defined as the number of pupae that emerged from the initial 24 hour egg lay.

### Microbiome profiling

The microbiomes were profiled using 16S rRNA amplicon sequencing. DNA was first extracted from pools (10 flies for each cage and time point) using the Quick-DNA Plus kit (Zymo D4068), which includes a proteinase K digestion to ensure unbiased sampling of diverse bacteria. Proteinase K digestion did not affect the characterization of the microbiome (Supp. Fig. M1), thus in our analysis, we computationally merged samples from the same sampling point with +/- proteinase K. 16S rRNA amplicons were generated using a two-step dual-indexed approach. We amplified the V1-V2 region of the 16S rRNA gene (Supp. Table M1), pooled for cleanup with Ampure XP beads, and then digested with BstZ17l enzyme to deplete *Wolbachia* amplicons (40). Libraries were sequenced using 300 bp paired-end reads using the Illumina MiSeq platform at the Princeton University Genomics Core.

Sequences were processed using QIIME2 v2020.6 (80). DADA2 was used to cluster the amplicon sequence variants (ASVs) (81). Taxonomy was assigned using the Greengenes reference database (82), trimmed to the 16S rRNA V1-V2 region. Phyloseq was used to visualize data (83). Potential contaminants were flagged using the decontam package (84) and removed prior to analyses. Samples were rarefied to 1000 reads per pool for analyses (Supp. Fig. M2 for rarefaction).

### Statistical analyses

To determine if *Wolbachia* altered the capacity for the microbiome to change during the seasonally fluctuating environment, we first calculated alpha-diversity. We calculated Shannon diversity and Faith’s phylogenetic diversity on ASVs. We then calculated the change for both diversity measures from the prior sampling point and summed the absolute value of change. We tested for significant differences between *Wolbachia* status using the Kruskall-Wallis test.

For beta-diversity, we first examined differences across all cages at all timepoints. We calculated Bray-Curtis dissimilarity (BC) for all samples and then used PERMANOVA implemented in vegan (85) to test for the effects of *Wolbachia* and time (over the course of the growing season) on community structure. To better understand how beta-diversity changed over time, we then examined beta-diversity change within each cage. We determined the change in BC with both the complete community and only the top four most abundant bacterial genera: *Acetobacter, Commensalibacter, Providencia,* and *Watuersellia;* four genera together comprised ~85% across all samples. The comparison between the complete community and the top four abundant bacteria allowed us to understand whether dominant microbes interact more with *Wolbachia* than low abundance bacteria. We used a mixed linear model to test for the effects of *Wolbachia,* time, and their interaction on BC, with cage as a random effect implement in lme4 in R (86).

To assess the phenotypic effects of *Wolbachia* and microbiome interactions, we used *Commensalibacter* as a covariate in the starvation resistance analyses. *Commensalibacter* was the most frequently occurring bacteria for the three phenotyping timepoints (Fig. 2). Given that the starvation resistance assay itself would alter the microbiome, we could not directly assess the microbiome of the flies we phenotyped. However, we used the *Commensalibacter* value from the preceding microbiome sample (i.e., Day 92 microbiome for the Day 96 phenotyping point). For starvation resistance, we fit a mixed effect Cox proportional hazard model for each timepoint separately. We modeled the response of starvation resistance (i.e., time to death) considering the effects of *Wolbachia,* sex, and *Commensalibacter* relative abundance and with cage as a random effect, implemented in the coxme package in R (87). We tested for *Wolbachia* x sex interactions, but if the interactions were non-significant, they were removed from the model.

To determine whether *Wolbachia* and the microbiome influenced fly fitness, we included the ratio of *Providencia* to *Commensalibacter* (Prov:Comm) as our microbiome covariate. We included the Prov:Comm ratio to more fully capture variation in the microbiome, rather than just a single bacteria in the model. Both were frequently found in flies, with *Providencia* in 6/8 cages and *Commensalibacter* in all eight. We modeled the response using a mixed-effect model with the negative binomial as the error distribution in the glmmTMB package (88). Our model tested for the effect of *Wolbachia,* lifespan, Prov:Comm interaction, and the three way interaction on fecundity, with cage as the random effect. We assessed significance of the main effects using Wald X^2^ tests with Kenward-Roger degrees of freedom.

## DATA AVAILABILITY

Amplicon sequencing will be uploaded to NCBI [upon acceptance]. Phenotyping data will be uploaded to Dryad [upon acceptance]. Code used to analyze data will be posted on github [upon acceptance].

## ACKNOWLEDGEMENTS

We thank Princeton Facilities for space and support in maintaining the field populations. B. Haseley, T. Price, L. Price-Henry, and L. Papp provided invaluable help in building the field cages. S. Rudman and P. Schmidt provided helpful feedback in sampling design. LPH was supported by NSF-GRFP under grant DGE1656466 and National Institutes of Health (NIH) grants GM124881 to JFA.

## SUPPLEMENTAL METHODS

### Construction of field mesocosms

The field site was located at Princeton University, NJ (40.34°N, 74.64°W). Cages were 1.2 m x 0.6 m x 0.6m (height x width x depth) and constructed from polyethylene monofilament fabric with 150×150 micron mesh (Greenhouse Megastore IS-NT-99) and spring locks (Greenhouse Megastore GF-9004). Eight total cages were constructed at the field site (Fig. 1). Within each cage, we provided shading for flies through a shelving unit and dwarf peach tree without fruit. Each cage held a temperature/humidity data logger (Elitech USA Temlog 20H). Fly food was composed of 10% glucose, 10% yeast, 1.2% agar, with 0.04% phosphoric acid and 0.4% propionic acid as preservatives. The fly food was provided in aluminum foil loaf pans (Web Restaurant Store #612A80).

### Phenotyping procedures

To understand how Wolbachia and microbiome change affected fly fitness, we collected age-matched flies for phenotyping. To age-match flies, we performed an additional egg lay in a 2.5” muffin tin (Reynolds) with ~80 ml of fly food. The muffin tin was placed inside a small cage within the larger cage, allowing flies to develop with the same environmental and microbial conditions as the rest of the population. Flies were age-matched to 5-7 days old, and then brought into the lab for phenotyping. Flies were collected on Day 96, Day 116, and at the end Day 127 for starvation resistance. We note Day 127 flies were not specifically age matched, but collected from within the larger cage, which reflects the standing variation in traits at the end of the season. For this reason, we primarily focus on differences within each sampling point, rather than comparing between the three sampling points.

To measure starvation resistance, individual flies were placed in custom acrylic arenas each 15mm in diameter and 6.25mm in height. Arenas only contained 1% agar to provide humidity, but no nutritional value. Individuals were allowed to freely move around their individual arenas. Flies were aspirated into the plates to avoid side effects from CO_2_ anesthetization on behavior (80). Cages were randomized across observation plates. After plating flies, we recorded their movements at 1 frame per second using Basler acA3088-57um cameras allowing for full frame videos at 3088×2064. This yields approximately 8.197 px/mm – sufficient resolution to robustly identify individuals and their movements. All recordings were taken with LoopBio’s Motif and compressed with libx264. After all flies died, we determined the time of death from the video recordings. To do this, we checked the videos every 2.5 hours to find the time when death occurred to quantify starvation resistance.

For the Day 127 flies, we also measured fecundity and lifespan from individual females to identify whether these fitness-related traits varied between Wolbachia positive and negative flies. Individual females were placed into fly vials with 6 ml fly food and allowed to lay eggs for ~24 hrs. Fecundity from females was measured as the number of pupae that emerged from the 24 hour egg lay. We then flipped the flies into new vials for 10 days. Then, after 10 days, we flipped every 3-4 days until flies died to determine the lifespan of each individual. We considered fecundity to be the number of pupae that emerged from the initial 24 hour egg lay.

**Supp. Fig. 1:**
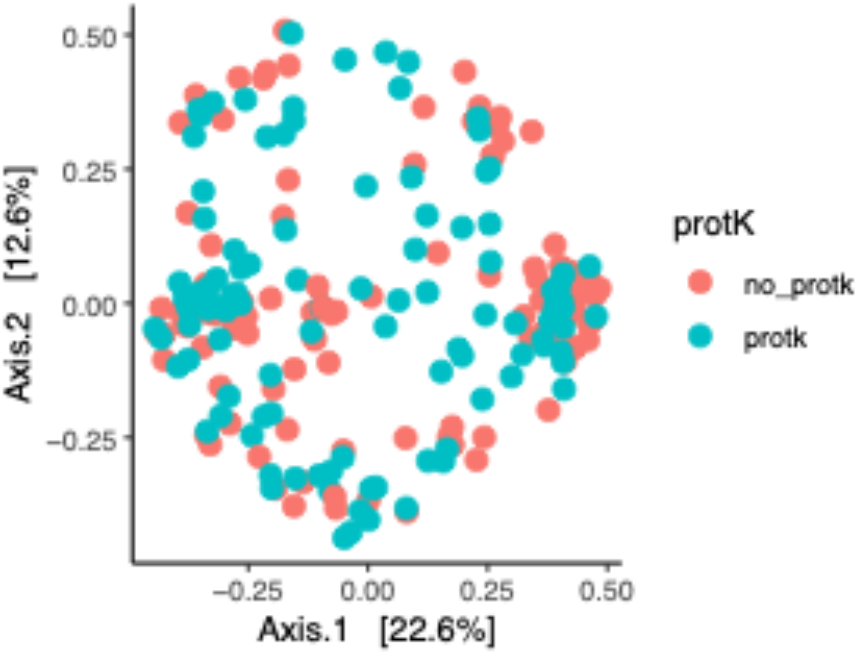
PCoA plot based on Bray-Curtis dissimilarity, colored by proteinase K treatment. Each point represents a sample, colored by proteinase K treatment. There was no statistically significant difference between proteinase K treatments (PERMANOVA, F_1,213_ = 1.7, R^2^ = 0.008, p=0.058).

**Supp. Fig. 2:**
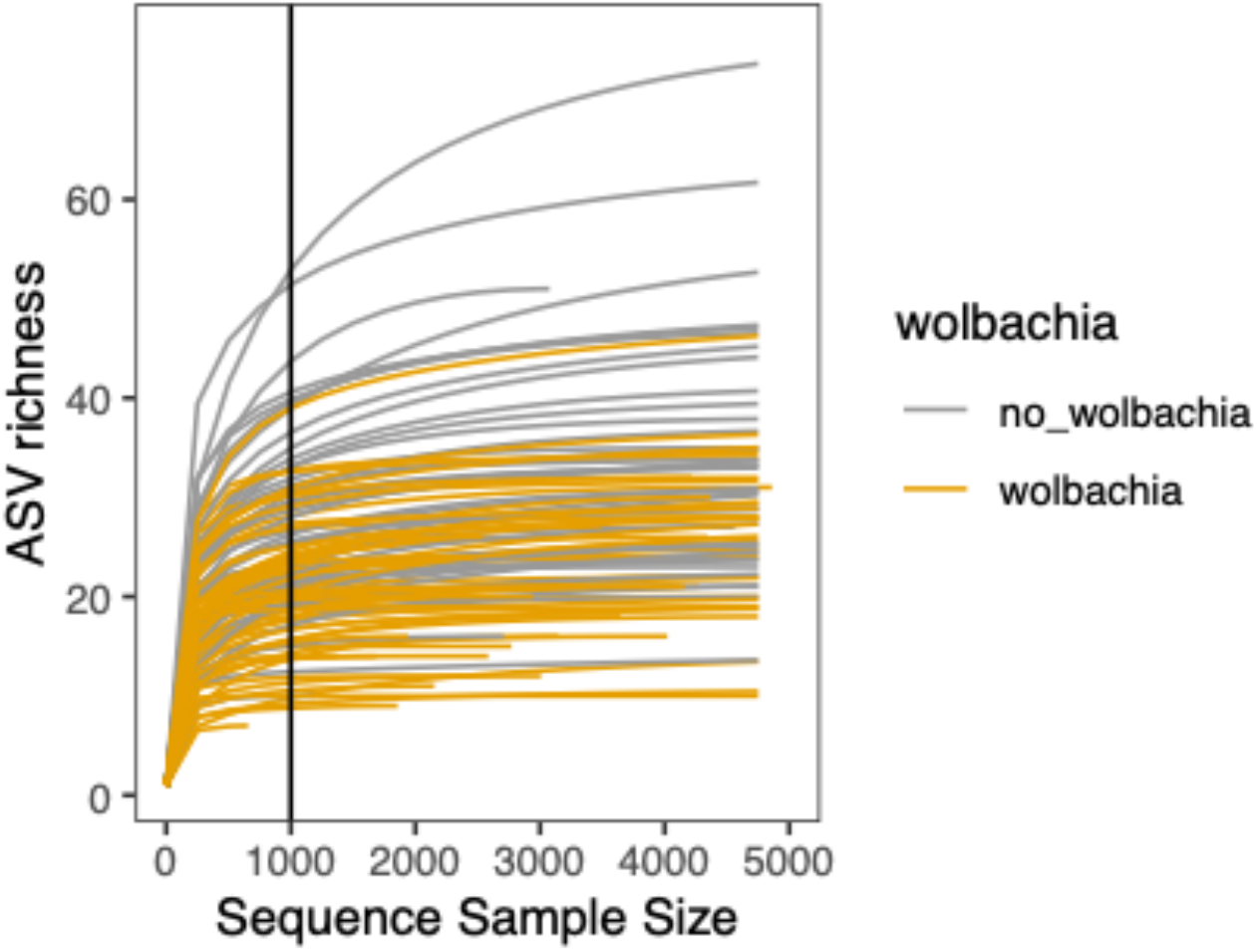
Rarefaction curves of microbiome samples. Each line represents a different sample, colored by *Wolbachia* status. The vertical line shows the rarefaction level (1000 reads/sample) used in the microbiome analyses presented here. In general, ASV richness was captured within 1000 reads/individual, though we do note a few samples had much higher ASV richness, and the 1000 reads/individual likely did not capture the ASV richness completely. However, the 1000 reads/sample allowed for us to maintain most samples.

**Supp. Table R1:** PERMANOVA: Bray-Curtis ~ Wolbachia + Time + (strata = cage) PERMANOVA results for Bray-Curtis dissimilarity between Wolbachia status and time (days since start of experiment). Asterisks denotes significance (i.e.: * p<0.05, ** p<0.005, *** p<0.001).

| **PERMANOVA: Bray-Curtis ~ Wolbachia + Time + (strata = cage)** |  |  |  |  |  |  |
| --- | --- | --- | --- | --- | --- | --- |
|  | **Df** | **Sums Of Squares** | **Mean Squares** | **F.Model** | **R<sup>2</sup>** | **Pr(>F)** |
| Wolbachia | 1 | 0.64 | 0.6396 | 2.3883 | 0.01805 | 0.001 *** |
| Time | 1 | 7.203 | 7.203 | 26.8956 | 0.20332 | 0.001 *** |
| Residuals | 103 | 27.585 | 0.2678 |  | 0.77863 |  |
| Total | 105 | 35.428 |  |  | 1 |  |

**Supp. Table R2:** PERMANOVA: Unifrac ~ Wolbachia + Time + (strata = cage) PERMANOVA results for Unifrac distance between Wolbachia status and time (days since start of experiment). Asterisks denote significance (i.e.: * p<0.05, ** p<0.005, *** p<0.001).

| **PERMANOVA: Unifrac ~ Wolbachia + Time + (strata = cage)** |  |  |  |  |  |  |
| --- | --- | --- | --- | --- | --- | --- |
|  | **Df** | **Sums Of Squares** | **Mean Squares** | **F.Model** | **R<sup>2</sup>** | **Pr(>F)** |
| Wolbachia | 1 | 0.8634 | 0.86342 | 6.7909 | 0.05362 | 0.027 * |
| Time | 1 | 2.1445 | 2.14447 | 16.8664 | 0.13317 | 0.001 *** |
| Residuals | 103 | 13.0959 | 0.12714 |  | 0.81322 |  |
| Total | 105 | 16.1038 |  |  | 1 |  |

**Supp. Table R3:** Model: Bray-Curtis (top 4 abundant bacteria) ~ Time * Wolbachia + (1|cage) Fixed effects for community turnover (mean Bray-Curtis dissimilarity by each population) over the growing season for the top four abundant bacteria. Significance was evaluated using Type III Wald F-tests with Kenward-Rogers degrees of freedom. Asterisks denotes significance (i.e.: * p<0.05, ** p<0.005, *** p<0.001).

| | Estimate | Std. Error | df | t value | Wald $X^2$ | Pr ($>X^2$)<sup>b</sup> |
| --- | --- | --- | --- | --- | --- | --- |
| (Intercept) | 0.8815 | 0.0211 | 28.0515 | 41.859 | 1752.1443 | < 2.2e-16 *** |
| Time | -0.0019 | 0.0002 | 92.4282 | -9.099 | 82.7876 | < 2.2e-16 *** |
| Wolbachia | -0.0506 | 0.0192 | 75.1030 | -2.641 | 6.9729 | 0.0083 ** |
| Time x Wolbachia | 0.0008 | 0.0002 | 91.9450 | 3.593 | 12.9091 | 0.0003 *** |

<sup>b</sup>df=1 for all variables

**Supp. Table R4:**
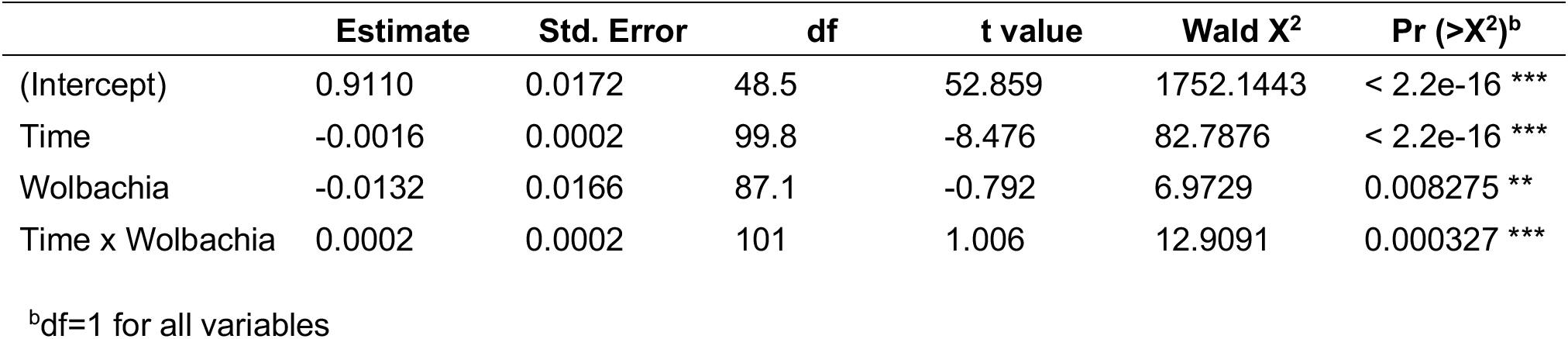
Model: Bray-Curtis (all bacteria) ~ Time * Wolbachia + (1|cage) Fixed effects for community turnover (mean Bray-Curtis dissimilarity by each population) over the growing season for the complete microbiome. Significance was evaluated using Type III Wald F tests with Kenward-Rogers degrees of freedom. Asterisks denotes significance (i.e.: * p<0.05, ** p<0.005,*** p<0.001).

**Supp. Table R5:**
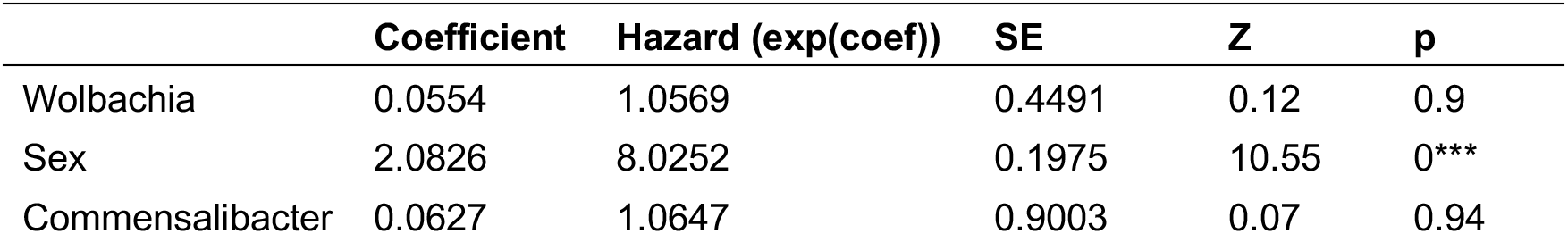
Model: Starvation time ~ Wolbachia + sex + Commensalibacter + (1|cage) Day 96 phenotyping. Effects of Wolbachia, sex, and Commensalibacter on starvation resistance using Cox mixed-effects model fit by maximum likelihood. Asterisks denote significance (i.e.: * p<0.05, ** p<0.005, *** p<0.001).

**Supp. Table R6:**
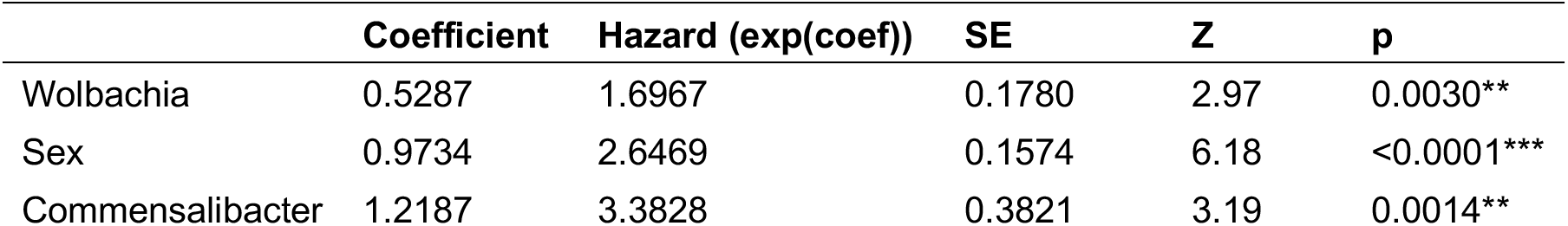
Model: Starvation time ~ Wolbachia + sex + Commensalibacter + (1|cage) Day 116 phenotyping. Effects of Wolbachia, sex, and Commensalibacter on starvation resistance using Cox mixed-effects model fit by maximum likelihood. Asterisks denote significance (i.e.: * p<0.05, ** p<0.005, *** p<0.001).

| **Model: Starvation time ~ Wolbachia + sex + Commensalibacter + (1\|cage)** |  |  |  |  |  |
| --- | --- | --- | --- | --- | --- |
|  | **Coefficient** | **Hazard (exp(coef))** | **SE** | **Z** | **p** |
| Wolbachia | 0.5287 | 1.6967 | 0.1780 | 2.97 | 0.0030** |
| Sex | 0.9734 | 2.6469 | 0.1574 | 6.18 | <0.0001*** |
| Commensalibacter | 1.2187 | 3.3828 | 0.3821 | 3.19 | 0.0014** |

**Supp. Table R7:** Model: Starvation time ~ Wolbachia + sex + (Wolb. * sex) + Commensalibacter + (1|cage) Day 127 phenotyping. Effects of Wolbachia, sex, Wolbachia * sex interaction, and Commensalibacter on starvation resistance using Cox mixed-effects model fit by maximum likelihood. Asterisks denote significance (i.e.: * p<0.05, ** p<0.005, *** p<0.001).

| **Model: Starvation time ~ Wolbachia + sex + (Wolb. \* sex) + Commensalibacter + (1\|cage)** |  |  |  |  |  |
| --- | --- | --- | --- | --- | --- |
|  | **Coefficient** | **Hazard (exp(coef))** | **SE** | **Z** | **p** |
| Wolbachia | -0.0004 | 0.9996 | 0.2722 | 0 | 1.0000 |
| Sex | 0.9075 | 2.4780 | 0.2241 | 4.05 | 0.0001*** |
| Commensalibacter | 1.3307 | 3.7838 | 0.4790 | 2.78 | 0.0055** |
| Wolbachia * sex | -0.7097 | 0.4918 | 0.2980 | -2.38 | 0.0170* |

**Supp. Table R8:** Model: Fecundity ~ Lifespan * Wolbachia * PC ratio + (1|cage) Fixed effects for the effects of lifespan, Wolbachia, microbiome (Providencia:Commensalibacter ratio) on fecundity using generalized linear mixed model with negative binomial as the error distribution. Asterisks denote significance (i.e.: * p<0.05, ** p<0.005, *** p<0.001).

| **Model: Fecundity ~ Lifespan \* Wolbachia \* PC ratio + (1\|cage)** |  |  |  |  |
| --- | --- | --- | --- | --- |
|  | **Estimate** | **Std. Error** | **Z** | **Pr(>\|Z\|)** |
| (Intercept) | 2.5838 | 0.3538 | 7.304 | <0.0001*** |
| Lifespan | 0.0049 | 0.0085 | 0.579 | 0.5629 |
| Wolbachia | 0.9914 | 0.3270 | 3.032 | 0.0024** |
| Prov:Comm (PC) ratio | 1.4700 | 0.6886 | 2.135 | 0.0328* |
| Lifespan*Wolbachia | -0.0150 | 0.0085 | -1.765 | 0.0776 |
| Lifespan*PC ratio | -0.0141 | 0.0178 | -0.794 | 0.4271 |
| Wolbachia*PC ratio | -2.6697 | 0.6912 | -3.863 | 0.0001*** |
| Lifespan*Wolbachia*PC ratio | 0.0444 | 0.0178 | 2.488 | 0.0128* |

**Supp. Table M1:**
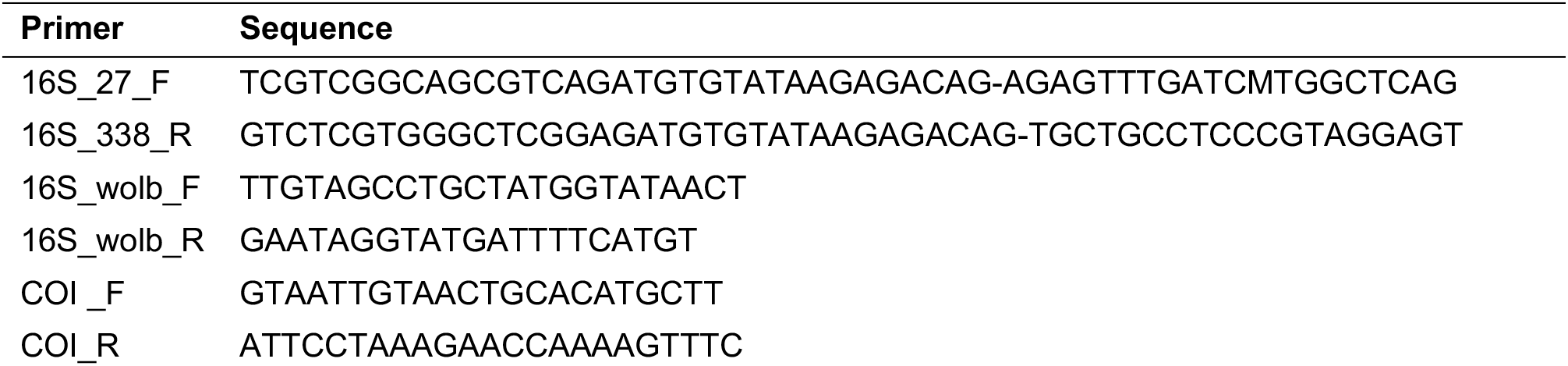
Primer sequences used. For the amplicon sequencing primers (16S_27_F and 16S_338_R), the dash separates the Illumina adapters from the 16S rRNA locus.

